# Chromatin Changes Associated with Neutrophil Extracellular Trap (NET) Formation in Whole Blood Reflect Complex Immune Signaling

**DOI:** 10.1101/2025.01.04.631302

**Authors:** Justin Cayford, Brandi Atteberry, Akanksha Singh-Taylor, Andrew Retter, Benjamin P. Berman, Theresa K Kelly

## Abstract

**Background:** Neutrophils are key players in innate immunity, forming neutrophil extracellular traps (NETs) to defend against infections. However, excess NET formation is implicated in inflammatory conditions such as sepsis and immunothrombosis. Studying NET formation in isolated neutrophils provides important mechanistic insights but does not reflect the complexity of immune interactions in whole blood, limiting our understanding of neutrophil responses.

**Methods:** This study investigates chromatin accessibility changes using Assay for Transposase-Accessible Chromatin with sequencing (ATAC-Seq) during phorbol 12-myristate 13-acetate (PMA) induced NET formation in whole blood. We compared chromatin accessibility patterns in neutrophils following PMA treatment in isolation and whole blood to assess the impact of other immune cells and signaling environment.

**Results:** Whole blood PMA stimulation elicited consistent chromatin accessibility changes across donors, demonstrating organized chromatin decondensation during NET formation. The chromatin response was characterized by increased accessibility in genomic regions enriched for immune-specific pathways, highlighting the role of immune cell interactions in NET formation. Differentially accessible regions (DARs) present following PMA induction in whole blood and isolated neutrophils showed greater association with NET-related and inflammatory transcription factors, while DARs specific to isolated neutrophils showed fewer relevant motifs. Pathway analysis indicated that whole blood responses involved more robust activation of immune-specific pathways, such as interleukin and cytokine signaling, compared to isolated neutrophils.

**Conclusions:** Our findings underscore the importance of studying NET formation within a whole blood environment to capture the complexity of neutrophil responses and immune cell interactions. This understanding is crucial for identifying effective therapeutic targets in NET-associated inflammatory diseases.

## 1. Introduction

Neutrophils are integral to the innate immune response, being among the first cells to respond to infection (1, 2). They detect pathogens through various cell surface receptors - activated directly by pathogens, or indirectly via interactions with other immune cells, such as platelets and monocytes (3, 4). Upon detection, neutrophils eliminate pathogens through phagocytosis, degranulation, and the release of NETs—web-like structures composed of chromatin, histones, and antimicrobial proteins that trap and neutralize pathogens (5, 6). Controlled NET formation benefits the host; however, dysregulated NET release leads to excessive inflammation, tissue damage, thromboinflammation, disseminated intravascular coagulation, and organ dysfunction (7, 8). Sepsis is associated with significant changes in neutrophil function, with the severity of dysregulation correlating with disease severity (9, 10). Interactions with other immune cells, including platelets and monocytes, are crucial in such disease states, significantly influencing neutrophil activation and NET formation (11).

Chromatin decondensation is a pivotal step in NET formation (12), preceding DNA expulsion from neutrophils. Various cellular mechanisms and transcription-related processes mediate this decondensation (13). ATAC-Seq is a powerful method for identifying open chromatin regions, making it ideal for assessing early chromatin structure changes during NET formation (14). Understanding the molecular mechanisms behind chromatin decondensation and NET formation can identify therapeutic targets to modulate maladaptive NET formation and has the potential to improve patient outcomes.

Current research on NETs utilizes neutrophil-like cells from immortalized cell lines (15), *in vitro* models with isolated primary neutrophils (16), and mouse models (17). While isolated neutrophil studies provide insights into direct stimuli effects, they cannot capture indirect effects mediated by other circulating cells. For instance, platelets play a crucial role in neutrophil activation and recruitment, contributing to NET formation (18), and platelet-neutrophil complexes regulate inflammatory feedback loops that lead to excessive NET release if uncontrolled (19). Macrophages and monocytes also influence NET formation through cytokine signaling and cellular interactions (20). Consequently, signaling mechanisms underlying NET formation may differ between isolated neutrophils and those in whole blood, where these interactions occur *in vivo*.

Neutrophil isolation methods influence the cell’s sensitivity and response to NET stimuli (21). Prolonged isolation procedures can induce stress in neutrophils, potentially altering their state and behavior (21). Thus, while isolated neutrophils offer a controlled environment to study NET formation, they do not fully reflect *in vivo* activity, as lack of external signaling that is present in whole blood may alter the neutrophil response. *In vivo* mouse studies maintain intact whole blood but differ significantly from the human immune system (22), limiting the translation potential of findings. These limitations highlight the need for innovative approaches to study NET formation in a more clinically relevant environment.

To better understand chromatin changes associated with NET induction and assess the impact of the human whole blood environment, we performed ATAC-Seq time courses to map chromatin accessibility changes in neutrophils following PMA treatment in whole blood. Previously, we demonstrated that PMA induction led to chromatin accessibility changes associated with NET formation in isolated neutrophils (23). By comparing NET formation in whole blood and isolated neutrophils, we identified genomic regions associated with neutrophil activation and NET formation in both environments, as well as regions specific to either isolated or whole blood induction. Chromatin accessibility changes that occurred in both isolated and whole blood systems reflected neutrophil processes like degranulation and regulation of innate immune responses. While whole blood-specific regions were enriched for more immune-specific pathways and cytokine signaling, highlighting the influence of the complex blood environment on neutrophil responses. Notably, chromatin accessibility changes upon PMA induction were more pronounced in the whole blood system compared to the isolated system, suggesting immune crosstalk in the whole blood environment could amplify the neutrophil response. These findings emphasize that chromatin decondensation during NET formation is influenced by interactions within whole blood, underscoring the importance of studying NET formation in a clinically relevant environment to identify molecular biomarkers and develop NET inhibitors.

## 2. Materials and Methods

### 2.1 Ethics Approval

Whole blood was obtained from healthy donors (Supplementary Table 1) in K2-EDTA tubes (BD #366643) (PrecisionMed, San Diego). Research was approved under WCG IRB Protocol number 20161665, conforming to the Declaration of Helsinki and the Human Belmont Report nd all participants provided written informed consent. Each subject was healthy, aged 18-50, with BMI < 30, and not taking NSAIDs.

### 2.2 Whole Blood Treatment

Whole blood was pooled, and 20 mL aliquots distributed into two 50 mL tubes (ThermoScientific, Cat#339653). Untreated samples were collected as 2 mL aliquots in 5 mL tubes (Eppendorf, Cat#0030119401) and fixed as described below. The remaining 18 mL was treated with either 250 nM PMA (Sigma, Cat#P1585) or DMSO (ATCC, Cat#4-X). Subsequently, 2 mL aliquots were placed in 5 mL tubes for each time point (30 min, 60 min, 90 min, and 120 min) and maintained at 37°C until further processing.

The cells were fixed with a 10x solution of formaldehyde (Formaldehyde 11% (Sigma, Cat#252549), 1M NaCl, 0.1 mM EDTA (FisherScientific, Cat#AM9010), 0.5mM HEPES (ThermoFisher, Cat#15630080), added at 1:10 volume to whole blood aliquots. After incubation at room temperature for 10 minutes, fixation was quenched with a 1:20 volume of glycine (2.5M) (ThermoFisher, Cat#15527013). Neutrophils were then isolated from the fixed whole blood using the MACSxpress Whole Blood Neutrophil Isolation Kit for humans (Miltenyl Biotec #130-104-434). Briefly, the bead mixture was added to the whole blood sample, followed by incubation on a rotator. Magnetic separation yielded isolated neutrophils, which were washed in 1x PBS and subjected to red blood cell (RBC) lysis to eliminate contaminants. Cells were washed in ice-cold 1x PBS and pelleted to retain only intact neutrophils, removing potential NET fragments or other DNA. 125,000 cells per condition/time point/replicate were flash frozen in liquid nitrogen and stored at –80°C until further processing.

### 2.3 FACS Analysis

Cell staining and FACS analysis were conducted at the La Jolla Institute of Immunology as previously described(24). Briefly fixed/frozen cell pellets were reconstituted in ice-cold 1x PBS. Staining was performed using PE-A: CD66b (BD Bioscience Cat # 561650) to identify the positive neutrophil population, APC-A: CD14 (BD Bioscience Cat # 555399) for the negative neutrophil population, and Alexa Fluor 488-A: CD45 (BD Bioscience Cat # 567402) and BV650-A: CD16 (BD Bioscience Cat # 563691) for additional characterizations. The obtained results were analyzed using FlowJo software to determine the percentage of purity for the isolated neutrophil populations according to the described methodologies.

### 2.4 Tn5 Assembly

Recombinant Tn5 transposase protein (Active Motif #81284) was assembled with custom oligos mosaic end (ME) ME_Rev, ME_A, and ME_B (IDT), and activity was tested as previously described (14).

### 2.5 ATAC-Seq Protocol

The fixed ATAC-Seq protocol was performed as previously reported (23). Briefly, adapted from Chen et al. (25), nuclei were transposed in a reaction mixture containing assembled Tn5 transposase, followed by cross-link reversal, DNA purification, library preparation, and sequencing.

### 2.6 ATAC-Seq Processing and Peak Calling

ATAC-Seq processing and alignment were conducted using an ATAC-Seq Nextflow pipeline (https://nf-co.re/atacseq/2.1.2) (26) with the nf-core framework as previously described (23). Samples were aligned to the hg38 reference genome, with the fragment size parameter set to 200. Peak calling was performed using MACS2 (27), identifying narrow peaks at a false discovery rate (FDR) of 0.01. This pipeline followed current ENCODE sequencing standards (28).

### 2.7 Untreated ATAC-Seq Analysis

Consensus peaks (.featureCounts.txt) and annotated peaks (.annotatePeaks.txt) were generated using the nf-core pipeline for subsequent analysis. Read counts were normalized across samples using scaling factors from bigwig normalization (https://cran.r-project.org/web/packages/scales/index.html) to balance total reads. For untreated samples, normalized peak counts were combined with annotated peaks to create a consensus peak dataset. UpSetR plots (https://cran.r-project.org/web/packages/UpSetR/index.html) were generated using the .boolean.annotatePeaks.txt files. Additional metrics, including read counts, peak annotations, bigwigs, fraction of reads in peaks (FRiP) scores, insert sizes, and alignment metrics, were generated using the nf-core ATAC-Seq pipeline (Supplementary Table 2).

### 2.8 Treated ATAC-Seq Analysis

Treated samples were analyzed similarly to untreated samples, with the following modifications. Samples were normalized and divided into six groups based on time course (T30, T60, T90, and T120 minutes) and treatment (DMSO vs. PMA). Samples with less than 10 million reads or a FRiP score below 5 were excluded (Supplementary Figure 1B-D). Peaks were retained if at least 2/3 of donors at a given timepoint/treatment had a peak within that interval. For DESeq2 (29) analysis, normalized counts of consensus peaks were filtered to include only those with a baseMean > 50, and subsequent analyses were performed on this dataset.

Principal Component Analysis (PCA) was conducted using prcomp, and pairwise comparisons were performed with DESeq(), summarizing results with the results() function. Significant regions (padj < 0.01) were z-score normalized based on normalized peak counts, and heatmaps were generated using pheatmap (https://cran.r-project.org/web/packages/pheatmap/index.html) with row clustering. Volcano plots were created using DESeq2’s results() function, defining significant regions by -log10(padj) > 4. For all-vs.-all comparisons, significant regions from each grouping (treatment and timepoint) with padj < 0.01 were included. Pheatmap (https://cran.r-project.org/web/packages/pheatmap/index.html) was used with both row and column clustering. Overlaps between T30 DMSO vs. PMA T30-T120 were analyzed using inner_join (tidyverse) (30) and visualized with eulerr version 7.0.2 for Venn diagrams in a custom R script.

### 2.9 HOMER Analysis

HOMER was used to determine motif enrichment in differentially accessible regions (DARs) after PMA stimulation, as previously described (23, 31), with the following modifications. Significant regions were sorted by padj, and a background set of DARs was randomly selected from the total consensus peaks generated in Nextflow for the dataset (.featureCounts.txt) for motif enrichment. Next, findMotifsGenome.pl was used with hg38 and standard settings and results were visualized using a custom R script.

### 2.10 Isolated Data Processing

Isolated neutrophil data was obtained from the Sequence Read Archive (SRA) (https://www.ncbi.nlm.nih.gov/sra, BioProject: PRJNA1120432) (23). This data was processed using the same pipeline and settings as untreated and treated ATAC-Seq analysis, including alignment, normalization, peak calling with MACS2, and DESeq2 analysis. Replicate-merged datasets for isolated samples were used in all comparisons.

### 2.11 Pathway Analysis

DAR gene lists from ATAC-Seq were analyzed for pathway enrichment using the following bioinformatics tools: DAVID (https://david.ncifcrf.gov) was used to identify functional domains and motifs. Gene lists were divided into whole blood and isolated neutrophil datasets, and functional annotation was performed using Reactome, KEGG, and GO databases to identify pathways relevant to NET formation. Metascape (https://metascape.org) (32) was also used to analyze the same gene lists. Enrichment analysis was conducted through Metascape’s express analysis mode, followed by custom downstream analysis using Python3. Results were processed using the Pandas library (v2.2.0)(26), focusing on the top 20 p-values per condition, with missing values replaced by 0. NumPy (v1.26.4)(26) handled missing values, and heatmaps were generated using Seaborn (v0.13.2)(33) with Matplotlib (v3.8.3) (34) for customization. These heatmaps visualized enrichment scores (-log10(pvalue)) with average clustering and significance annotations. EnrichR gene set enrichment analysis (https://maayanlab.cloud/Enrichr/) (35) was used to compare two gene sets. Each gene set was uploaded separately to the tool to identify enriched pathways (configured to focus on Reactome pathways). For each dataset, pathway enrichment results, including adjusted p-values and z-scores, were retrieved to highlight significant pathways. Overlapping pathways between the two gene sets were identified to evaluate shared mechanisms. Unique pathways were also noted to highlight distinct regulatory networks in each gene set. Results were exported for downstream analysis.

The Reactome enrichment data was subsequently utilized in Python3 for figure generation. Custom scripts were written to process the enrichment data, focusing on p-values and pathway names. Visualization libraries such as Matplotlib (v3.8.3) and Seaborn (v0.13.2) were employed to create publication-quality figures, showcasing key pathways and their significance. The final figure highlighted both shared and unique pathways, providing a clear comparative analysis of the two gene sets.

## 3. Results

### 3.1 Whole Blood Fixation Followed by Neutrophil Isolation Shows Stable Chromatin Structure Across Donors

Previous work has assessed chromatin changes associated with NET formation in isolated neutrophils (23, 36-38). To expand upon this and understand the interplay of the whole blood matrix and potential immune interactions, we induced NET formation in whole blood, followed by formaldehyde fixation, neutrophil isolation, and ATAC-Seq. Whole blood was either untreated (n=6), PMA stimulated (n=5), or DMSO stimulated (n=5) (Figure 1A, left), and compared to neutrophil ATAC-Seq data where cells were isolated prior to fixation (21) (Figure 1A, right). To minimize the impact of stress during isolation, we fixed whole blood with formaldehyde prior to neutrophil isolation, aiming to maintain neutrophils close to their circulating state. This approach yielded neutrophil preparations of about 90% purity with monocytes as the main contaminant (Supplementary Figure 1A).

**Figure 1.**
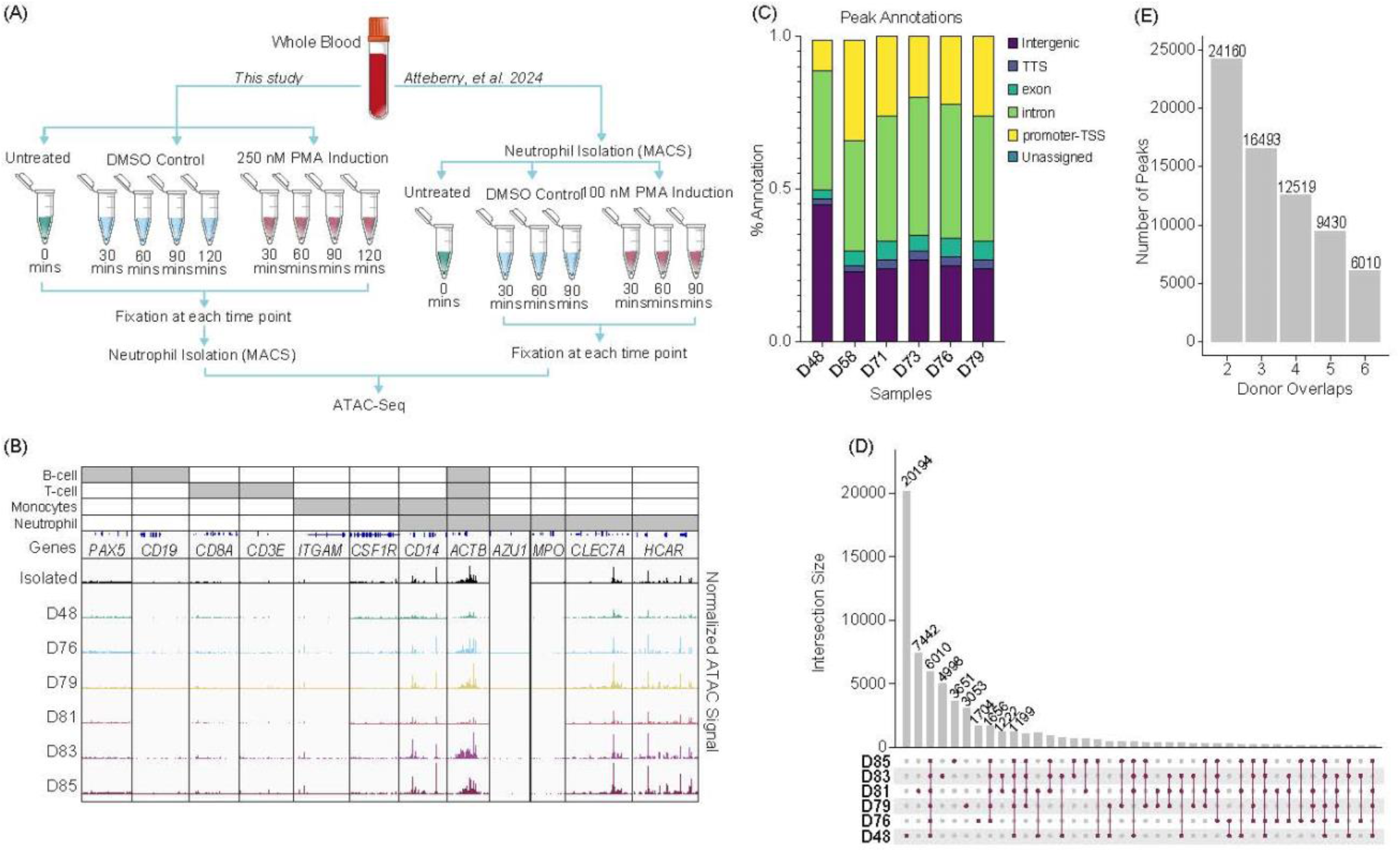
Neutrophils fixed in whole blood prior to isolation shows consistency across donors: **(A)** Experimental schematic. Left - Whole blood was collected, formaldehyde fixed and then isolated using the MACSexpress Whole Blood Neutrophil Isolation Kit for humans (Miltenyi Biotec #130-104-434). The whole blood was either untreated (n=6), stimulated with 250 nM phorbol 12-myristate 13-acetate (PMA) or dimethyl sulfoxide (DMSO) for the given time, fixed, isolated, followed by ATAC-Seq. Right – Data from isolated neutrophils was previously described and is shown here for clarity (23). **(B)** Merged replicate tracks visualized using the IGV Genome Browser with untreated healthy donors (n=6). Neutrophils isolated prior to fixation (top) (23) and whole blood fixed prior to isolation (D48, D78, D79, D81, D83, D85) are shown below. Top bars (grey) indicate loci specific to different immune cells: Housekeeping (all) – *ACTB*; B-cell (*PAX5* and *CD19*), T-cell (*CD8A* and *CD3E*), monocytes (*ITGAM, CSF1R*, and *CD14*), and neutrophils (accessible regions – *CD14, CLEC7A*, and *HCAR*; inaccessible regions – *AZU1* and *MPO*). **(C)** The location of peaks (MACS2 peaks (q < 0.01)) across genome structures was generated using nf-core/atacseq. Peaks annotations are plotted as a percentage of all peaks found within a sample (Transcription start site (TSS), Transcript termination sites (TTS)). **(D)** UpSetR plot for untreated whole blood fixed samples shows the peak intersections across donors. **(E)** The number of peaks that are shared across a given number of n=6 donors.

Untreated whole blood samples fixed prior to isolation (donors D48, D76, D79, D81, D83, and D85) showed high correlation across donors and with post-isolation samples (Isolated), with minimal presence of accessible chromatin which would be expected from contamination of other common immune cells (Figure 1B). We examined gene loci expected to have accessible chromatin in neutrophils (*CD14, ACTB, CLEC7A*, and *HCAR*) and regions expected to lack accessibility (*AZU1* and *MPO*) (23, 39). Despite lower purity in whole blood fixed samples, loci specific to B cells (*PAX5* and *CD19*), T cells (*CD8A* and *CD3E*), and monocytes (*ITGAM, CSF1R*) (40) showed no chromatin accessibility, as expected (Figure 1B). We also found accessibility in CD16/*FUT4*, but not CD33/*FCGR3B* and variability across donors in accessibility of CD15 (Supplementary Figure 1E). This suggests that chromatin structure in the whole blood model is consistent across donors and with what is observed in neutrophils that are isolated prior to fixation.

Called peaks were reproducible across donors and showed limited variability in gene annotations (Figure 1C). Specifically, 6,010 peaks were shared among all six donors, indicating a set of commonly accessible chromatin regions in whole blood neutrophils (Figure 1D). Although donors D48 and D81 had more unique peaks, the majority of peaks were shared across multiple donors, demonstrating considerable overlap.

Notably, over 24,000 peaks were found in at least two donors, while nearly 16,500 peaks were common across three donors (Figure 1E). This consistency highlights the robustness of chromatin accessibility profiles in whole blood neutrophils across donors, despite individual variability.

### 3.2 PMA Stimulation Drives a Dynamic Chromatin Response Over Time Compared to DMSO Controls

To assess chromatin accessibility changes during PMA-induced NET formation in whole blood, we treated whole blood from five donors (D48, D76, D79, D81, and D85) with 250 nM PMA or DMSO and selected time points prior to NET release (24). Samples were fixed at 30, 60, 90, and 120 minutes, followed by neutrophil isolation and ATAC-Seq. Samples with fewer than 10 million reads and a FRiP score below 5 were excluded (Supplementary Figure 2A, 2B). Called peaks (MACs, q < 0.01) and their annotations showed limited changes based on treatment conditions, with differences mainly due to donor variability (Supplementary Figure 2C, 2D).

To determine whether PMA treatment drove global changes in chromatin accessibility, we performed Principal Component Analysis (PCA) and observed separation between PMA and DMSO groups (Figure 2A). Comparing chromatin accessibility at various loci showed stability of *TBP* (housekeeping gene) and increased accessibility at *CXCL2, CXCL3, CXCL5*, and *CD69* loci in PMA-treated samples at 60 minutes (Figure 2B). A bimodal response at *ACTG1* indicated rapid opening followed by decreased accessibility by 90 minutes consistent with previous findings (23, 41). There were also regions with decreased accessibility, such as the *CXCR2* locus (Figure 2B). These results indicate a gain and loss of chromatin accessibility with PMA induction.

**Figure 2.**
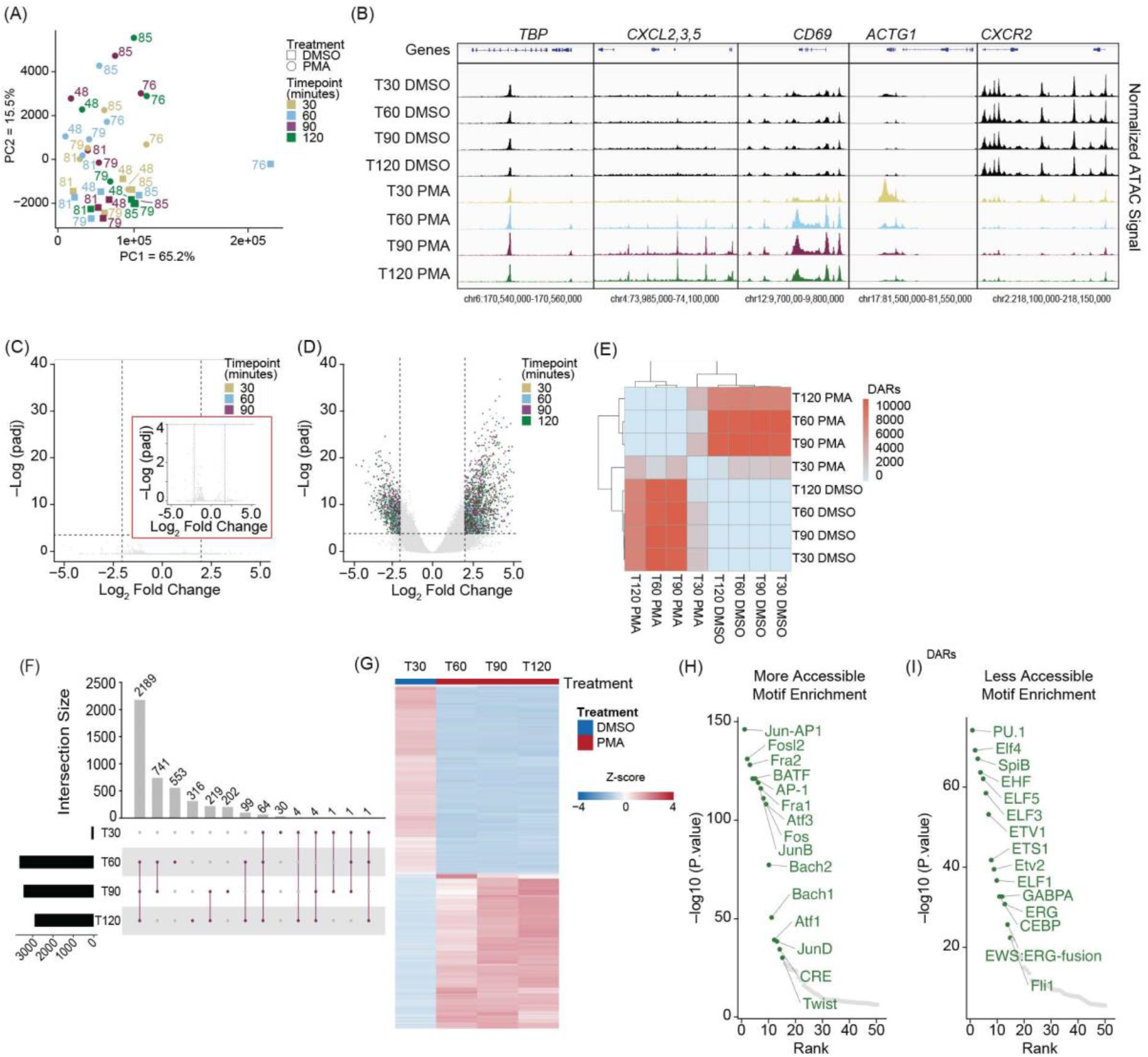
PMA stimulation drives a stable chromatin response in whole blood fixed neutrophils: **(A)** Principal Component Analysis (PCA) based on unbiased clustering of dimethyl sulfoxide (DMSO) versus phorbol 480 12-myristate 13-acetate (PMA) using the merged replicate data. Timepoints are indicated by color (T30 – gold, T60 – blue, T90 – purple, T120 - green). Treatment is indicated by squares (DMSO) or circles (PMA). PC1 (x-axis) represents 65.2% of the total variance and PC2 (y-axis) represents 15.5% of variance. Donor numbers are indicated by the number adjacent to the datapoint. **(B)** Merged donor tracks visualized using IGV genome browser. T30, T60, T90, and T120 DMSO (black), T30 PMA (gold), T60 PMA (blue), T90 PMA (purple), T120 PMA (green) at various loci. Housekeeping gene *TBP*, increased accessibility at the *CXCL2, 3, 5* locus (T60-T120), bimodal response shown at *ACTG1*, and decreased accessibility at *CXCR2*. **(C)** Volcano plot comparing T30 DMSO with T60, T90, and T120 DMSO. DESeq2 was used for a pairwise comparison and then plotted. The -log10(p.adj) value is graphed on the y-axis and Log2(fold change) is indicated on the x-axis. Inset graph (red outline) is zoomed in at the y-axis from 0-4 (-log(padj)). Dashed lines indicated significance thresholds (log2(Fold change) > 2.5 or < -2.5 and -log10(p.adj) > 4). There were no significant values (-log10(p.adj) > 4). **(D)** Similar to Figure 2B but comparing T30 DMSO vs. T30, T60, T90, and T120 PMA. Significant values (-log10(p.adj) > 4) were separated by timepoint. T30 – yellow, T60 – blue, T90 – purple, T120 – green. **(E)** Heatmap showing the total number of differential accessible regions (DARs) between all pairwise comparisons (each timepoint for each treatment condition) (DESeq2 p.adj > 0.01 and log2(fold change) less than -1.5 or greater than 1.5). **(F)** UpSetR plot of the differentially accessible regions (DARs) at each timepoint versus T30 DMSO (T30, 60, 90, 120 PMA). **(G)** Z-score normalized count heatmap of the top 1,000 DARs sorted on DESeq2 p.adj value for T30 DMSO versus PMA at T60, T90, and T120. Treatment is indicated by DMSO (blue) and PMA (red). Columns are organized by treatment and time. Rows are hierarchically clustered. **(H)** HOMER motifs within DARs that gain accessibility in T60-T1120 PMA compared to T30 DMSO are plotted by by-log10(p.value) on the y-axis and the HOMER rank on the x-axis. The top 50 known motifs were graphed, and the top 15 known motifs based on p.value are annotated in green. **(I)** Similar to (H), but DARs that have more accessibility in T30 DMSO compared to T60-T120 PMA are shown.

These results support the concept that PMA treatment induces chromatin changes preceding NET release. DESeq2 analysis revealed few significant DARs in the DMSO group between 30 and 120 minutes (Figure 2C), indicating chromatin stability across the time course. However, comparing DMSO at 30 minutes to PMA at all timepoints showed numerous significant DARs (Figure 2D, Supplementary Figures 2E-2H). Pairwise comparison of the number of DARs (q < 0.01) revealed that DARs were consistent and reproducible across the dataset, with limited chromatin changes at 30 minutes in PMA treatment, but significant changes at 60, 90, and 120 minutes compared to DMSO at 30 minutes (Figure 2E).

To gain a better understanding of the pattern of chromatin accessibility changes following PMA treatment, we determined the number of DARs across different time points. Consistent with Figure 2D, we found limited chromatin accessibility changes at 30 minutes, with only 105 DARs identified. Of these, 64 were present across all four time points (30-, 60-, 90-, and 120 minutes), and the majority of the changes occurred after 30 minutes. Specifically, 2,189 DARs were shared across the 60, 90 and 120 minutes of PMA (Figure 2F). Thus, the 30-minute PMA-treated timepoint was excluded from further analyses. The remaining PMA treated DARs (60, 90, 120 minutes) were combined to generate a comprehensive list and using DMSO at 30 minutes as a baseline, we assessed chromatin accessibility progression over time in response to PMA treatment. DARs were sorted by adjusted p-values, and the top 1,000 were z-score normalized (Figure 2G). We next determined whether certain transcription factor (TF) binding sites were enriched in regions that were gaining or losing accessibility and found that regions gaining accessibility had TF motifs associated with NET formation, including the AP-1 complex (JUN, FOS, ATF) (42) and FRA1/2 (Figure 2H). In contrast, regions losing accessibility were linked to differentiation and proliferation pathways, including the ELF family and PU.1 (Figure 2I).

### 3.3 PMA-Induced Chromatin Accessibility Changes in Whole Blood and Isolated Neutrophils Are Similar but Distinct

Comparing NET induction in isolated neutrophils (23) and whole blood, we first assessed if the order of isolation and fixation affected consensus peaks (defined as peaks shared by at least 50% of donors in each group) (21). Approximately 57% (14,420) peaks were shared between conditions (Figure 3A), with 8% (2,073 peaks) unique to whole blood fixed samples and 35% (8,813 peaks) unique to neutrophils fixed after isolation (Figure 3A). Overlapping peaks indicate a strong consensus between datasets and numerous peaks that occur only when fixation occurs after neutrophil isolation.

**Figure 3.**
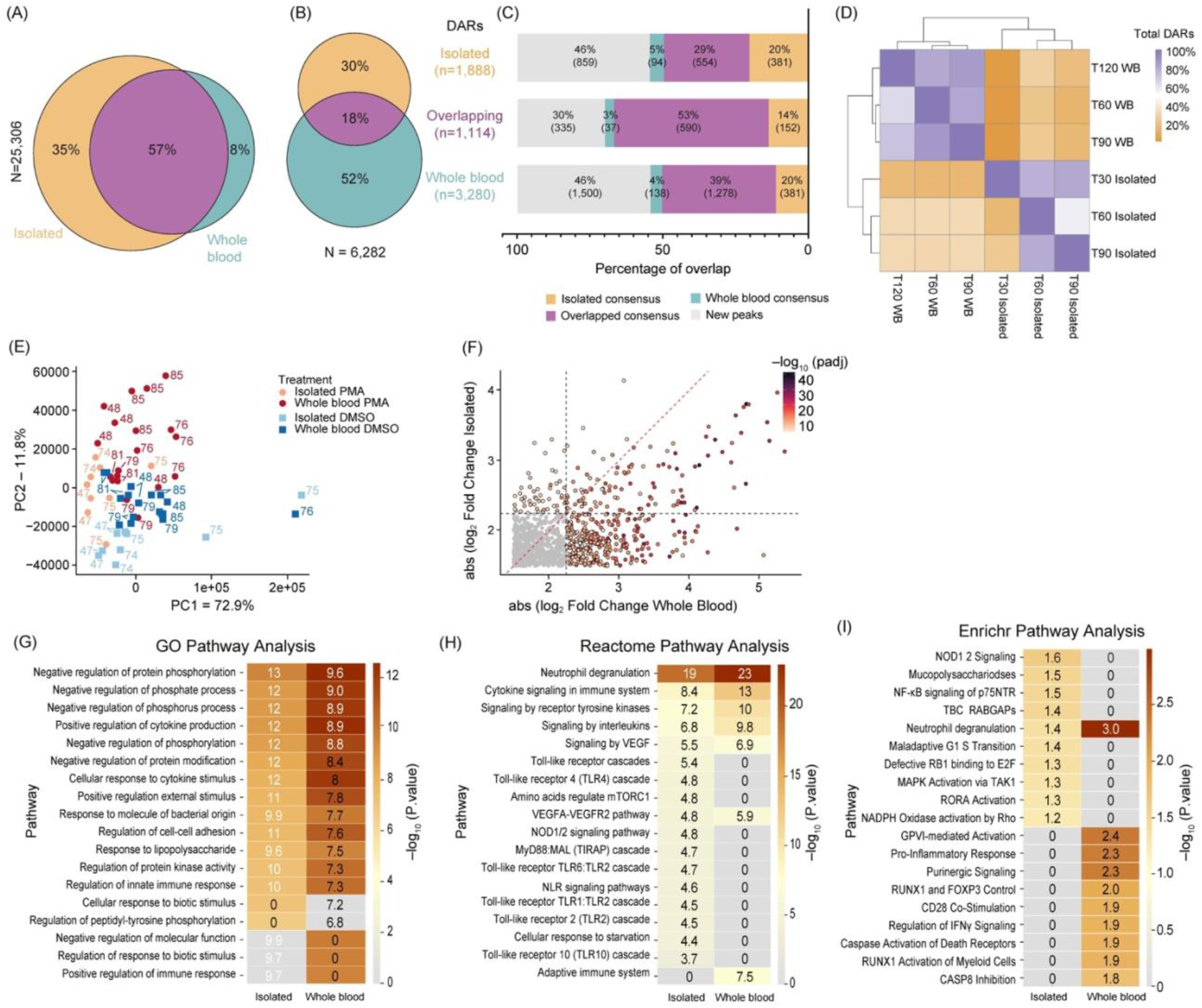
Comparison of chromatin accessibility changes between isolated and whole blood fixed neutrophils: **(A)** Venn Diagram showing overlaps between consensus peaks of untreated neutrophils (Isolated – orange; Whole blood – teal; Overlapping - purple) (minimum 50% of donors, n=6). Total peaks were 25,306, with percentages for unique and overlapping peaks indicated. **(B)** Venn Diagram of the DARs (DESeq2 q < 0.01) from Isolated (orange), Whole blood (teal), and Overlapping (purple). The total number of DARs is 6,282. **(C)** Stack bar plot showing the DARs separated into Isolated only (top, n=1,888), Overlapping (middle, n=1, 114), or Whole blood only (bottom, n=3,280). Bedtools intersect was used to determine the presence of DARs within the consensus peaks from (A). Categories include Isolated Consensus (orange), Overlapping Consensus (purple), Whole Blood Consensus (teal), and New Peaks (gray). Percentages and total DARs are listed within each bar. **(D)** Heatmap showing the overlap of DARs at each timepoint and condition. This indicates whether the DAR was also found at other timepoints between Isolated and whole blood (WB). The diagonal represents 100% of DARs called at each row, and subsequent values within the row represent the percentage of those DARs found. **(E)** Principal Component Analysis (PCA) clustering of Dimethyl sulfoxide (DMSO) versus phorbol 12-myristate 13-acetate (PMA) using merged replicate data. Datasets are indicated by color (Isolated PMA – light red, Isolated DMSO – light blue, Whole blood PMA – dark red, and Whole blood DMSO – dark blue). Treatment is indicated by squares (DMSO) or circles (PMA). PC1 (x-axis) represents 72.9% total variance and PC2 (y-axis) represents 11.8%. Donors are indicated adjacent to the datapoint. **(F)** Plotting the absolute value of DESeq2 log2(foldchange) values of all Overlapping DARs (n=1,114) between Isolated (y-axis) and Whole blood (x-axis). DARs were colored based on the lowest DESeq2 p.adj value between Isolated and Whole blood. Red dashed line indicates y=x and grey dashed lines indicate the Log2(FoldChange) threshold used for significant DARs (2.25). **(G)** GO pathway analysis of all DARs in Isolated and Whole blood. Each group includes Overlapping DARs. **(H)** Similar to (G) but Reactome pathway analysis was used. **(I)** Similar to (G) but Enrichr pathway analysis was used.

To determine whether NET associated chromatin changes are different when NETs are formed in isolated neutrophils or in the context of whole blood, we analyzed DARs following PMA induction in both systems. We identified 6,282 significant DARs, with 18% (1,114/6,282) overlapping between systems, 30% (1,888/6,282) unique to isolated neutrophils, and 52% (3,280/6,282) unique to the whole blood system (Figure 3B). PMA induced DARs were then assessed as to whether they were found in the consensus peaks from untreated isolated or whole blood neutrophils (Figure 3C). About 46% of PMA induced DARs were outside consensus peak sets, indicating PMA-induced new chromatin accessibility (Figure 3C). Interestingly, 70% of overlapping PMA induced DARs were found within original consensus peaks, compared to 60% in isolated and 54% of whole blood DARs (Figure 3C).

Temporal analysis of PMA-induced DARs showed consistent patterns across induction systems (Figure 3D, Supplementary Figure 3A). Despite only around 20% of the PMA induced DARs overlapping, samples generally clustered by treatment (DMSO vs. PMA) (Figure 3E). The DARs present in both isolated and whole systems exhibited stronger fold changes in whole blood compared to isolated samples (Figure 3F), with few divergent changes, indicating a robust PMA response across systems (Supplementary Figure 3B).

Gene Ontology (GO) analysis of DAR-associated genes revealed stronger enrichment in immune response pathways in whole blood compared to isolated samples (Figure 3G). Metascapes Reactome pathway analysis showed immune-specific pathways, including cytokine signaling and innate immune responses, were more prominent in whole blood (Figure 3H), whereas signaling cascades downstream from cell surface receptors and genes involved in response to starvation were more prominent in isolated neutrophils. EnrichR geneset enrichment analysis (35) revealed that the only category represented in both isolated and whole blood systems was neutrophil degranulation, with specific signaling pathways enriched in either isolated or whole blood systems (Figure 3I). These results suggest NET formation in whole blood involves more complex cell signaling pathways during PMA stimulation, reflecting interactions with other immune cells or factors present in whole blood.

### 3.4 Whole blood PMA induction leads to a more complex immune response than isolated neutrophil PMA induction

We next sought to further understand the role of the extracellular environment in NET formation by assessing changes between T30 DMSO and PMA treatments in both systems by z-score normalizing the DAR counts (n = 6,282) and mapping them over time (Figure 4A, Supplementary Table 3). DARs were next categorized based on changes in accessibility with PMA induction into isolated neutrophils only, whole blood only, or in both (i.e., overlapping) and the top half were retained (based on adjusted p-value) in each category for z-score normalization (Figure 4B). We observed that the isolated-only group had fewer DARs with increased accessibility, whereas whole blood and overlapping groups showed more balance of increasing and decreasing accessibility with PMA treatment (Supplementary Table 4).

**Figure 4.**
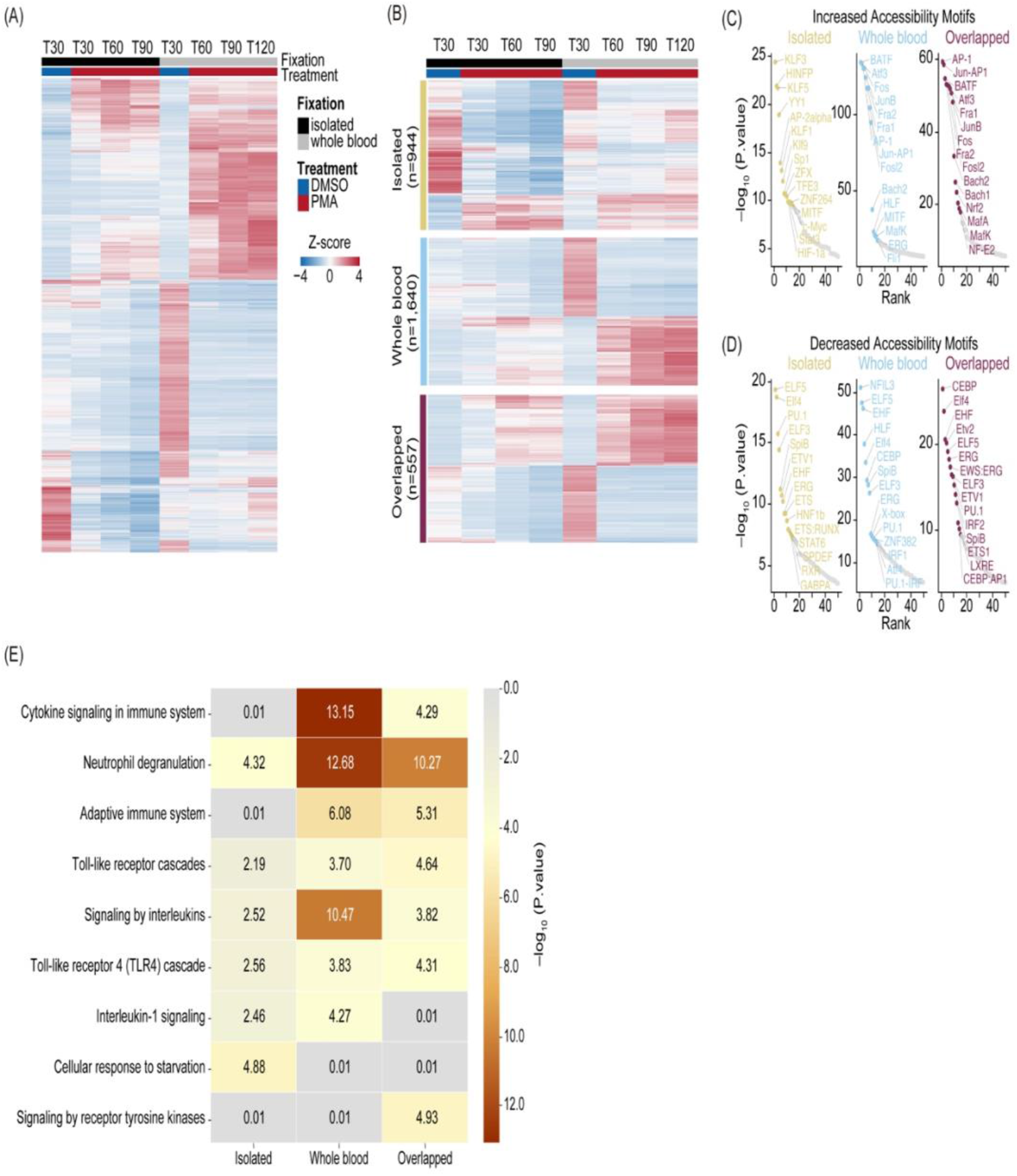
Whole blood PMA induction leads to a more complex immune response compared to the isolated system: **(A)** Heatmap of Z-score normalized count for the top DARs (sorted by p.adj) from Isolated, Whole blood, and Overlapped (n=6,282). T30 DMSO versus PMA at T30 (Isolated - black), T60, T90, and T120 (Whole blood - grey). Treatment is indicated by DMSO (blue) and PMA (red). Columns are organized by treatment and time. Rows were hierarchically clustered and represent different peaks. **(B)** Heatmap of Z-score normalized count of the top half DARs in each of the categories sorted on DESeq2 p.adj: Isolated (yellow – top; n=944), Whole blood (blue – middle; n=1,640), and Overlapped (purple – bottom; n=557). Treatment is indicated by DMSO (blue), and PMA (red) and fixation is indicated by Isolated (black) and Whole blood (grey). Columns are organized by treatment and time. Rows were hierarchically clustered. **(C)** HOMER motifs graphed by - log10(p.value) on the y-axis and HOMER rank on the x-axis for DARs which gained accessibility in Isolated (left – yellow), Whole blood (middle – blue), and Overlapped (right – purple). The top 50 known motifs were graphed, and the top 15 known motifs based on p.value are annotated. **(D)** Similar to (C) but with downregulated DARs that lose accessibility. **(E)** Reactome Pathway heatmap showing the -Log10(P.value) for each pathway: Isolated (left), Whole blood (middle), and Overlapped (right) DARs.

To better understand the underlying biology of NET formation in whole blood and isolated neutrophil environments, we separated DARs into increasing/decreasing chromatin accessibility and identified enriched TF binding motifs (Supplementary Table 5). We next identified overrepresented TF binding site sequences and found regions with increased accessibility in overlapping and whole blood groups were enriched for TFs associated with NET formation or inflammation, such as the AP-1 complex (43-45), and MAF/NFE2L2 (NRF2) (46) (Figure 4C). In the isolated group, the most identified motifs were not correlated with a NET response, except for HIF-1α and STAT3, which are known to play roles in neutrophil and inflammatory responses (Figure 4C) (47-49). These motifs were also present in whole blood and overlapping groups but were ranked below the top 20 based on p-value. Overall, DARs with decreased accessibility showed limited correlation to TFs directly involved in NET formation across all categories (Figure 4D).

We examined the genes associated with altered chromatin accessibility to identify affected pathways in response to PMA stimulation. Using Metascape’s Reactome pathway analysis (50, 51), we identified significant pathways in the three groups, finding many related to immune system regulation, including cytokine signaling, interleukins, and neutrophil degranulation. Interestingly, the top categories varied among whole blood, isolated, and overlapping groups. Immune pathways were less significant in the isolated group, likely because the main neutrophil response originated from overlapping regions in isolated samples (Figure 4E). In contrast, whole blood DARs showed much higher significance in interleukin and cytokine signaling compared to isolated or overlapping DARs (Figure 4E). Taken together, these data demonstrate dynamic changes in chromatin regions involved in NET formation, suggesting that similar yet distinct pathways govern NET formation in whole blood and isolated environments.

## 4. Discussion

Neutrophils are critical to the innate immune response, with NET formation being essential for responding to infections and modulating immune responses. However, maladaptive NET formation can lead to autoimmune diseases and a dysregulated host immune response such as seen in sepsis and acute respiratory distress failure (ARDS) (52-54). Given the complex blood milieu, understanding neutrophil induction and NET formation in the context of immune cell crosstalk is crucial. Chromatin accessibility is key in gene regulation, and investigating neutrophil responses under pathophysiological conditions can provide insights into maladaptive NET formation.

This study investigated differential chromatin accessibility in neutrophils within whole blood and compared them to isolated neutrophils post-PMA stimulation. Neutrophil isolation methods, whether through Ficoll gradients or commercial kits like MACSxpress, can significantly impact neutrophils (21) with the procedures potentially inducing stress responses and biasing experimental outcomes. The impact of the cell isolation procedures was shown through GO analysis which showed enrichment of genes associated with biotic stimulus response, and Reactome pathway analysis identified cell starvation, underscoring how isolation could alter neutrophil quiescence. Thus, studying neutrophils in a physiologically relevant environment is essential. Comparing whole blood to isolated neutrophils revealed NET formation requires higher PMA concentrations in whole blood and took longer following induction (24). While isolated neutrophils showed dramatic chromatin changes 30 minutes following PMA exposure, whole blood samples treated with PMA for 30 minutes looked more similar to DMSO treated samples with more dramatic chromatin changes happening at 60 minutes suggesting plasma components and cellular interactions may buffer or inhibit early NET formation. Due to the time it takes for neutrophil isolation the elapsed time from blood collection to fixation was more similar between the isolated 30-minute PMA and whole blood 60 minute PMA following timepoints so the contribution of elapsed time cannot be excluded. However, we note that there were limited chromatin changes with time in our DMSO treated controls. Despite timing differences, ATAC-Seq revealed dynamic chromatin accessibility changes, including increased, decreased, and bimodal responses.

Motif analysis identified TFs consistent with NET formation in both the whole blood and overlapping DARs, such as the AP-1 complex (43, 44), but not in the isolated only DARs. This indicates the whole blood *ex vivo* system may more effectively model NET formation by circulating neutrophils.

Fold change revealed a stronger chromatin response in whole blood samples as compared to isolated neutrophils (Figure 3F). The upregulation of cytokine/interleukin signaling pathways in whole blood suggests immune cell crosstalk, whereas isolated DARs showed limited activation of immune-specific pathways.

Enriched TF motifs in whole blood and overlapping categories were more NET-related, while isolated categories showed motifs unrelated to NET formation, except for HIF-1α and STAT3, which are involved in neutrophil and inflammatory responses (55). Furthermore, across all groups, motifs found in less accessible regions upon stimulation were associated primarily with TFs in the ELF family, PU.1, and ERG which are known to be involved in autoimmune disease (56-58). This suggests overlapped regions represent baseline neutrophil responses to PMA, while whole blood samples exhibit additional responses driven by the blood environment.

Our study demonstrates that examining NET formation within a human whole blood environment captures layered complexities of neutrophil responses by revealing additional chromatin accessibility changes and enriched immune-specific pathways not observed in isolated neutrophils. These changes likely reflect significant crosstalk among immune cells influencing neutrophil behavior during NET formation, pivotal in NET-associated pathogenesis such as immunothrombosis and sepsis.

Given the complex nature of human blood and immune responses—with diverse cellular components and myriad cytokines—studying NET formation within this intricate environment is critical for translating scientific findings into clinical applications. Our results indicate that strong synthetic stimulants like PMA lead to variable responses depending on the environment. Future work should focus on studying NET formation in whole blood using naturally occurring NET inducers, which are likely more subtle and may involve direct neutrophil actions or indirect signaling through cells like platelets and monocytes. Regulation of NET release by various cytokines and immune cell interplay will impact immune responses outcomes. Thus, a deeper understanding of the NETosis pathway in humans, considering these complexities and chromatin dynamics, is crucial for developing targeted therapies to modulate NETosis in inflammatory and autoimmune conditions.

## Supporting information

Supolementary Figures

## Data availability statement

The original contributions presented in the study are publicly available. This data can be found here: http://www.ncbi.nlm.nih.gov/bioproject/1177467. BioProject ID PRJNA1177467.

## Author Contributions

JC, TKK and AR conceived the study, JC, TKK, and BA designed the experiments. BA performed the experiments, JC and AST provided guidance, and BA, JC and BPB performed data interpretation. All authors participated in interpretation of data. BA, JC and TKK wrote the manuscript, and all authors edited the manuscript for important intellectual content.

## Acknowledgments

We would like to thank Christina Wheeler, Kieran Zukas and Finley Serneo, as well as the broader scientific team at Volition for helpful review of the data and discussion. ChatGPT version 4o was used to support the writing of R and Python3 codes and edits throughout the manuscript. We would also like to thank Cheryl Kim and Denise Hinz at the La Jolla Institute for Immunology for their FACS sorting work.

