## Supplementary material for "Chromatin Changes Associated with Neutrophil Extracellular Trap (NET) Formation in Whole Blood Reflect Complex Immune Signaling": Supolementary Figures

### 1 Figures

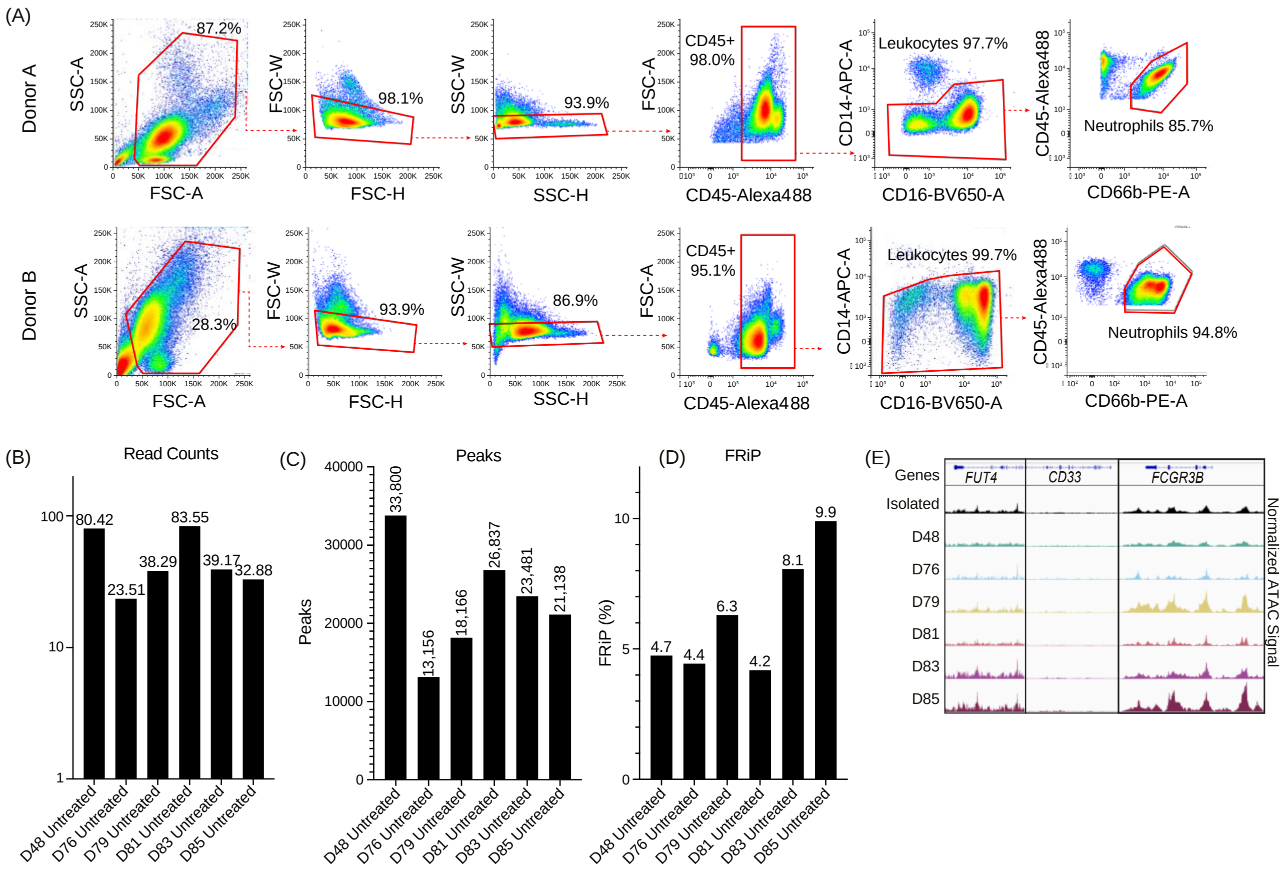


**Supplementary Figure 1: (A)** FACS results of neutrophils fixed in whole blood prior to isolation and gated through FSC-A vs SSC-A followed by FSC-H vs FSC-W and SSC-H vs SSC-W. CD45+ cells were selected and further separated to obtain CD16+CD14+ leukocytes. Monocytes were removed based on CD45+CD66b- and CD45+CD66b+ confirmed high neutrophil purity (~85.7% in Donor A and ~94.8%) in Donor B. **(B)** Total number of merged mapped reads for each untreated donor. The donors are indicated on the x-axis, and the number of mapped reads (in millions) is on the y-axis**. (C)** Similar to (B) but FRiP percent is shown on the y-axis. **(D)** Similar to (B) but total number of MACS2 peaks (q < 0.01). **(E)** Merged replicate tracks visualized using the IGV Genome Browser with untreated healthy donors (n=6). Neutrophils isolated prior to fixation (top – "Isolated") and whole blood fixed prior to isolated (D48, D78, D79, D81, D83, D85) are shown below. *CD15* (*FUT4*), *CD33*, and *CD16* (*FCGR3B*) are shown.

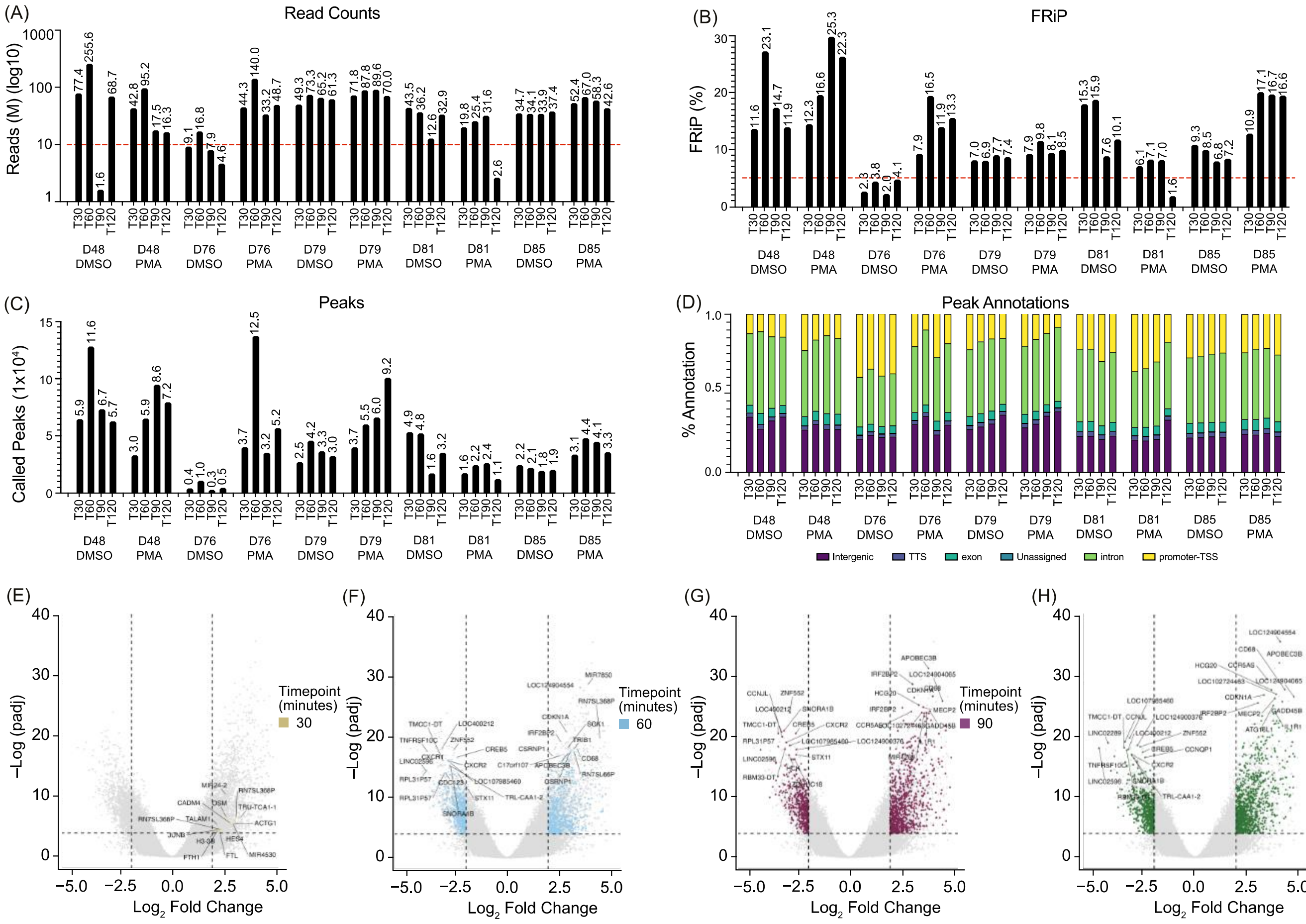


**Supplementary Figure 2: (A)** Total number of merged mapped reads for each of the donors, treatment, and time (x-axis). Total number of mapped reads (in millions) is on the y-axis. Dashed red line (y=10) indicates sequencing depth threshold. **(B)** Similar to (A) but FRiP percent is shown on the y-axis. Dashed red line (y=5) indicates FRiP threshold requirement. **(C)** Similar to (A) but with total number of MACS2 (q < 0.01) peaks. **(D)** Peak annotations generated on called MACS2 peaks (from (C)) for the samples shown as a percentage. (Transcription start site (TSS) Transcript termination sites (TTS)). **(E)** Volcano Plot comparing T30 DMSO with T30 PMA whole blood samples. DESeq2 was used for a pairwise comparison and then plotted. The -log10(p.adj) value is graphed on the y-axis and Log2(fold change) is indicated on the x-axis. Vertical dashed lines indicate thresholds at +/- 2.5 log2(Fold Change) and the horizontal dash line indicates threshold at -log10(p.adj) > 4. Grey points indicate regions that were not significant (abs(Log2(fold change)) > 2.5 and -log10(p.adj) > 4. Yellow color indicates significance and the top 15 positive and negative regions are labeled with the proximal gene to the differential peak. **(F)** Similar to Supplementary Figure 2E, but the comparison for significance was between T60 DMSO and T60 PMA. Blue indicates significant regions. **(G)** Similar to Supplementary Figure 2E, but the comparison for significance was between T60 DMSO and T60 PMA. Blue indicates significant regions. **(H)** Similar to Supplementary Figure 2E, but the comparison for significance was between T90 DMSO and T90 PMA. Maroon indicates significant regions. **(I)** Similar to Supplementary Figure 2E, but the comparison for significance was between T120 DMSO and T120 PMA. Green indicates significant regions.

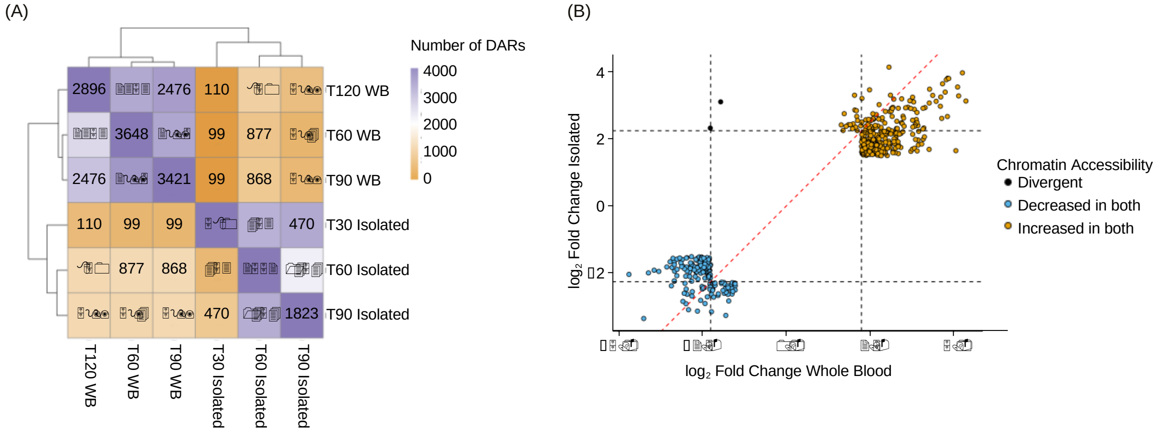


**Supplementary Figure 3: (A)** Heatmap showing the overlap of DARs at each timepoint and condition. This indicates whether the DAR was also found at other timepoints between Isolated and whole blood (WB). The diagonal represents the total number of DARs called at each row, and subsequent values within the row represent the total number overlapping of those DARs found. **(B)** Comparison of Log2(Fold change) for Isolated (y-axis) and Whole blood from DESeq2 DARs. Black dashed lines indicate fold change threshold (±2.25) and only DARs with values beyond these thresholds are shown in the graph. Red dashed line indicates y=x. DARs that were more accessible in both isolated and whole blood are shown in orange and DARs which were less accessible in both are shown in blue. Divergent DARs are shown in black.

### 2 Tables

<u>Supplementary Tables.xlsx</u>: (Table 1) Donor information for all donors in the study. (Table 2) Illumina Sequencing statistics of all samples. (Table 3) All 6,282 Differentially Accessible Regions (DARs) for both the Whole Blood and Isolated datasets (Table 4) Total number of DARs in each category (Isolated, Whole Blood, and Overlapping). (Table 5) HOMER Motifs generated for each of the comparisons.
